# Roadkill islands: carnivore extinction shifts seasonal use of roadside carrion by generalist avian scavenger

**DOI:** 10.1101/2021.02.18.429855

**Authors:** Matthew W. Fielding, Jessie C. Buettel, Barry W. Brook, Dejan Stojanovic, Luke A. Yates

**Affiliations:** School of Natural Sciences, University of Tasmania, Sandy Bay 7001 TAS Australia; ARC Centre of Excellence for Australian Biodiversity and Heritage; Fenner School of Environment and Society, Australian National University, Canberra, Australia

**Keywords:** road ecology, scavenging, trophic cascade, carcass use, Bayesian modelling, one-dimensional point patterns

## Abstract

1. Global road networks facilitate habitat modification and are integral to human expansion. Many animals, particularly scavengers, use roads as they provide a reliable source of food, such as carrion left after vehicle collisions. Tasmania is often cited as the ‘roadkill capital of Australia’, with the isolated offshore islands in the Bass Strait experiencing similar, if not higher, levels of roadkill. However, native mammalian predators on the islands are extirpated, meaning the remaining scavengers are likely to experience lower interference competition.
2. In this study, we use a naturally occurring experiment to examine how the loss of mammalian carnivores within a community impacts roadside foraging behaviour by avian scavengers.
3. We monitored the locations of roadkill and forest ravens (*Corvus tasmanicus*), an abundant scavenger species, on eight road transects across the Tasmanian mainland (high scavenging competition) and the Bass Strait islands (low scavenging competition). We represented raven observations as one-dimensional point patterns, using hierarchical Bayesian models to investigate the dependence of raven spatial intensity on habitat, season, distance to roadkill and route location.
4. We found that roadkill carcasses were a strong predictor of raven presence along road networks. The effect of roadkill was amplified on roads on the Bass Strait islands, where roadside carrion was a predictor of raven presence across the entire year. In contrast, ravens were more often associated with roadkill on Tasmanian mainland roads in the autumn, when other resources were low. This suggests that in the absence of competing mammalian scavengers, ravens choose to feed on roadside carrion throughout the year, even in seasons when other resources are available. This low interference competition could be disproportionately benefiting forest ravens, leading to augmented raven populations and changes to the vertebrate community structure.
5. Our study provides evidence that scavengers modify their behaviour in response to reduced scavenger species diversity, potentially triggering trophic shifts and highlighting the importance of conserving or reintroducing carnivores within ecosystems.

## Introduction

Land-use change across the world is facilitated and accompanied by road networks, which are expected to continue expanding rapidly over the next century (Laurance et al. 2014). For many animal species, expansive road networks have a number of negative impacts, such as vehicle collisions, fragmentation and dispersal barriers (Fahrig & Rytwinski 2009; Benitez-Lopez et al. 2010). However, some animals, particularly scavengers, appear to benefit from roads (Lambertucci et al. 2009; Planillo et al. 2015). Carrion is sporadic in wild environments and can often be difficult to locate, therefore, scavengers may regularly forage near roads because they provide a reliable and accessible source of carrion (Forman & Alexander 1998; Morelli et al. 2014). Tasmania, a large island (64,519 km^2^) south of Australia, is often regarded as the ‘roadkill capital of Australia’ (Hobday & Minstrell 2008; Nguyen et al. 2019), however, the smaller Bass Strait islands situated north of Tasmania could arguably lay an even greater claim to this title.

The Bass Strait islands were once part of a land bridge that connected Tasmania to the Australian Mainland, before sea-level rise flooded the basin ~ 14,000 years ago (Bowdler 2015). The productive and fertile land of the two largest islands in the region, King Island (1,098 km^2^) and Flinders Island (1,367 km^2^), have been heavily modified for agriculture with almost two-thirds of the native vegetation on King removed for farming (Threatened Species Section 2012). This has led to an upsurge of wallabies (*Thylogale billardierii* and *Macropus rufogriseus*) on the islands, with a recent census estimating half a million macropods on King Island (Branson 2008). Due to this high density, roadkill is a common sight, and farmers will often shoot the macropods *en masse* under crop-protection permits, leaving the bodies where they fall (Threatened Species Section 2012). Furthermore, land-use change on the islands has led to the recent (19^th^ – 20^th^ Century) extirpation of mammalian predators, such as quolls (*Dasyurus maculatus* and *Dasyurus viverrinus*), and the loss of the wedge-tailed eagle (*Aquila audax*) on King Island (Peacock et al. 2018; Fielding, Buettel & Brook 2020).

The extinction of carnivores within an ecosystem can have cascading effects throughout the food web (Allen et al. 2014; Ripple et al. 2014). Scavenging is an essential ecological function for maintaining stability in the ecosystem, via the control of community composition and carrion availability (Wilson & Wolkovich 2011; Barton et al. 2013; Buechley & Şekercioğlu 2016). As most carnivores are facultative scavengers, their loss can influence herbivore density and prompt mesopredator release (O’Bryan et al. 2019). For example, in north-east Tasmania, where Tasmanian devils (*Sarcophilus harrisii*) have declined due to devil-facial-tumour disease, feral cats (*Felis catus*) and forest ravens (*Corvus tasmanicus*) have increased in abundance due to increased access to carrion (Cunningham et al. 2018). In the Bass Strait, the surviving meso-predators could have greater access to carrion, leading to increased abundance and unbalance within the community structure relative to that elsewhere in Tasmania.

The forest raven is a common scavenger species present on both mainland Tasmania and the Bass Strait Islands (Higgins et al. 2006; Lawrence 2009). This species is known to feed on roadkill and, when in high abundance, could negatively impact other bird species through competition and direct killing (Debus & Rose 2006; Talmage 2011; Fielding, Buettel, Nguyen, et al. 2020). The species can also impact farming practices, attacking stock and raiding crop fields (Rowley 1969; Rowley & Vestjens 1973). Previous research found that forest ravens were the major beneficiary of increased carrion availability from the disease-driven population decline of devils (Cunningham et al. 2018). However, it is unknown how the complete loss of native mammalian predators may impact the scavenging behaviour of the species seasonally and on roadsides.

In this study, we recorded the locations of roadkill and forest ravens on eight road transects across the Bass Strait islands and Tasmanian mainland. This case study, with its mensurative experimental control (i.e., mammalian predators being present on mainland roads, but ‘removed’ from island roads) is ideal for understanding how the foraging behaviour of a generalist scavenger varies under lower levels of competition, and provides insights into the consequences of trophic shifts, highlighting the importance of conserving a suite of vertebrate scavenger species within a community. By representing raven observations as one-dimensional point patterns and using hierarchical Bayesian models, we investigate in a spatially explicit way how the dependence on roadside carrion by a generalist scavenger varies across habitat, season, distance to roadkill and areas with contrasting scavenger diversity. Point-pattern methods have been underutilised in one-dimensional data analysis, and the novel Bayesian methodology developed within this study has broad applicability to other transect data within ecological research.

## Methods

### Data Collection

Study sites were located in south-eastern Tasmania and the Bass Strait region (Fig. 1). Eight transects were selected, two within each of the Huon Valley, Tasman Peninsula, King Island and Flinders Island areas (see Fig. 1 map) based on the following criteria: a total length of 15 km, a speed limit of 80 km/h or above, two lanes of traffic, avoidance of urban areas, and the presence of both roadkill and live animals. On the eight transects, we collected presence and location data for roadkill and forest ravens. Road surveys were completed September 2016 – July 2017 for the south-eastern Tasmanian routes, and July 2019 – May 2020 for the Bass Strait roads. Within each season, each route was repeat surveyed four times with the repeats combined to give a set of observations for each season on each route (i.e., 4 seasons × 8 routes = 32 replicates). Survey-route direction was also alternated to minimise observer bias. Surveys were completed during daylight hours with each survey within a season being completed at different times of the day to minimise temporal bias in diurnal range. Roadkill and forest-raven-location data were recorded by driving at 40–50 km/hr and geolocating (Garmin eTrex 30) all roadkill and forest raven observations. Any forest raven which was observed on or adjacent to the road was recorded.

**Figure 1:**
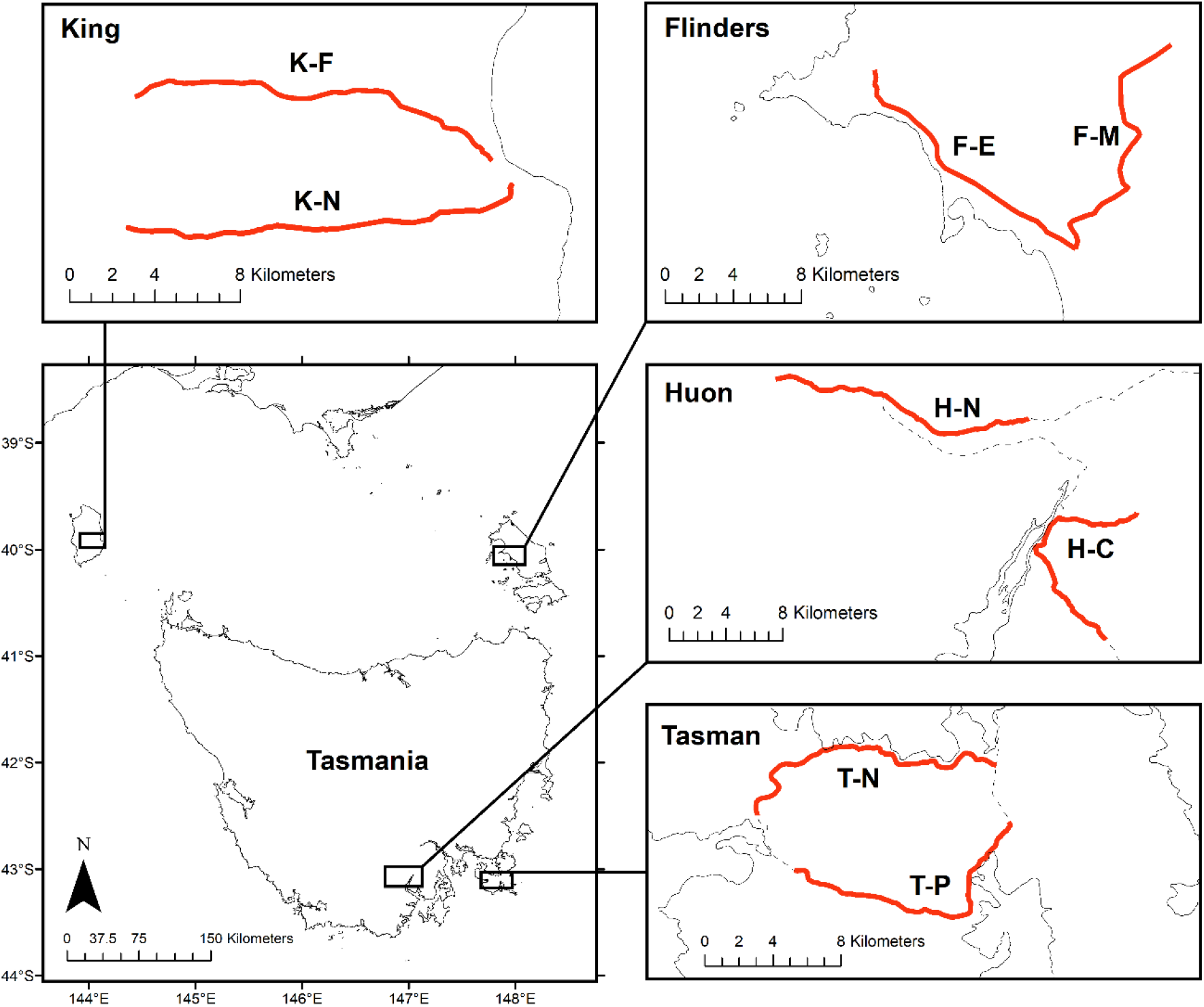
Geographic location of the survey routes in South-eastern Tasmania and the Bass Strait Islands.

### Statistical Analysis

Raven observations were represented as points on a linear network, resulting in 32 unique linear point patterns (pairs of season and route combinations; Fig. 2a). The distribution of points in a spatial point pattern is typically modelled as a Poisson process, which assumes that each observation is independent (Baddeley et al. 2020). Roadkill locations and habitat type were used to define spatial co-variates, *X_F_*: distance in metres to nearest open farmland; and *X_R_*: distance in metres to nearest roadkill (Fig. 2b). We defined a log-linear model of the spatially heterogeneous Poisson intensity *λ* = *λ*(*x*, *y*):

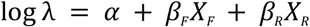

corresponding to the multiplicative model,

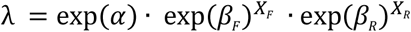

such that the (exponentiated) intercept, exp(*α*), gives the expected density at zero distance from farmland and zero distance from a roadkill.

**Figure 2.**
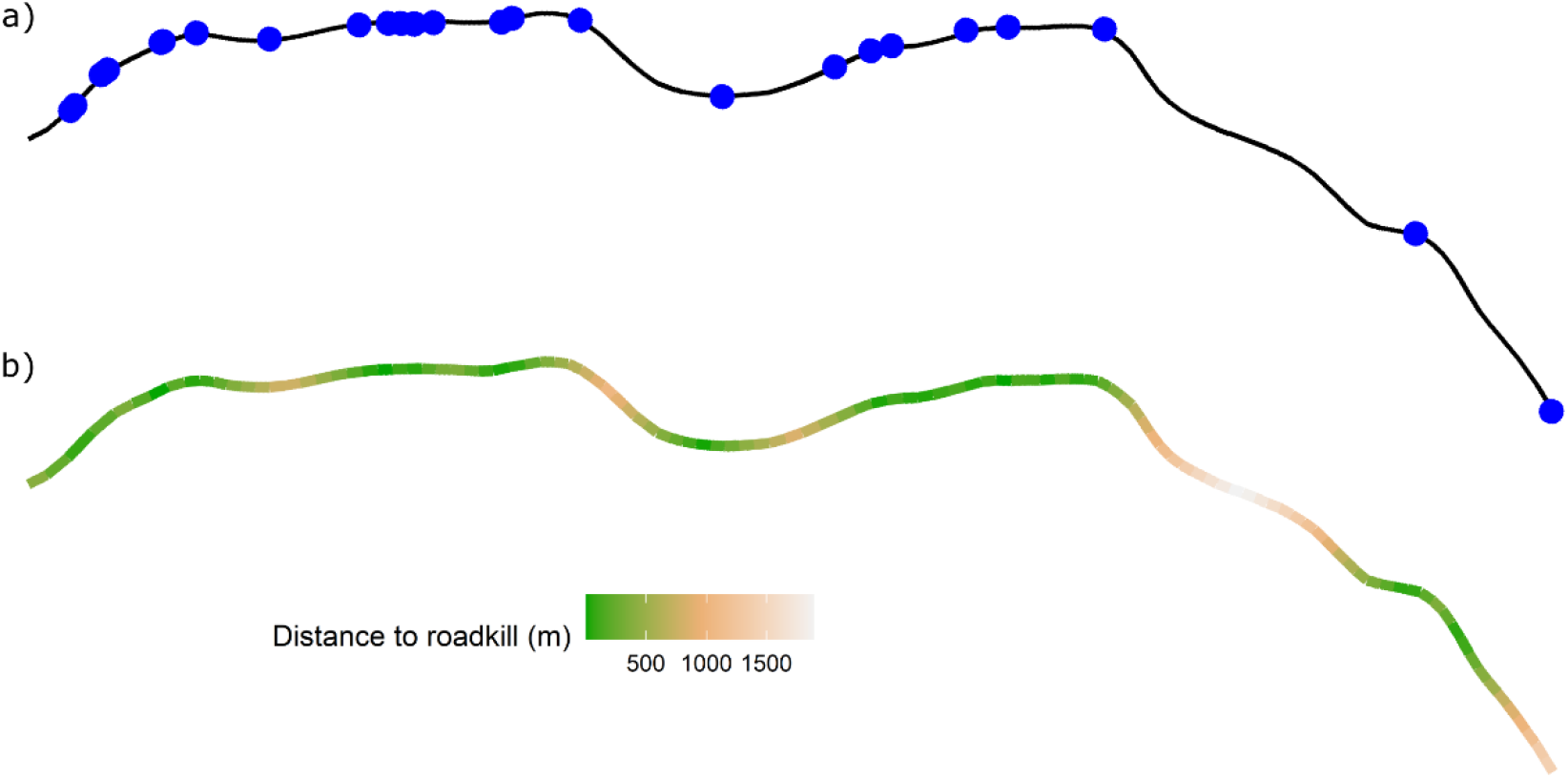
a) Raven observations from the K-F route in the season of spring represented as points on a linear network; b) Distance in metres to nearest roadkill on the K-F route in spring.

The model likelihood was approximated using a Berman-Turner device (Berman & Turner 1992), which encodes the point process as a numerical quadrature scheme, with spatial observations viewed as spatially discrete, Poisson-distributed events. We constructed each scheme using the R package *spatstat* (Baddeley et al. 2015), applying the default settings to generate approximately 15,000 quadrats for each of the 15 km routes. The 32 schemes were combined into a single data set with grouping variables for *season* and *route*.

To investigate the effects of both season and location with controlling for non-independence of repeats on a given route, we defined a set of candidate models characterised by different structural implementations of the corresponding fixed and random effects. The use of random effects maximised the utility of our data, permitting the inclusion of certain route-season pairs with fewer observations, thus retaining data for every route in every season. Model variants included: pooled vs unpooled grouping; diagonal vs full covariance matrices; treating *route* as a fixed vs random effect; and nested vs crossed effects (Table 1).

**Table 1.** Set of candidate models to investigate the effects of season and location on raven spatial intensity along roadsides. The model set contain several varied conditions including pooled vs unpooled grouping; diagonal vs full covariance matrices; treating *route* as a fixed vs random effect; and nested vs crossed effects. p_eff_ is the effective degrees of freedom (complexity) value, computed from the cross-validation estimate. N () and MVN () are normal and multivariate normal distributions, respectively.

| Form ($\log \lambda_{s,r}$) | Hierarchy | Group effect | Covariance matrix | $p_{\text{eff}}$ |
| --- | --- | --- | --- | --- |
| 1 $\beta_0$ | - | - | - | 1 |
| 2 $\beta_0 + \beta_F X_F + \beta_R X_R$ | - | Pooled | - | 3 |
| 3 $\beta_{0,s} + \beta_{F,s} X_F + \beta_{R,s} X_R$ | - | Fixed | - | 12 |
| 4 $\beta_{0,s} + \beta_{0,r} + (\beta_{F,s} + \beta_{F,r}) X_F + (\beta_{R,s} + \beta_{R,r}) X_R$ | - | Fixed / Crossed | - | 33 |
| 5 $\beta_{0,sr} + \beta_{F,sr} X_F + \beta_{R,sr} X_R$ | - | Unpooled | - | 96 |
| 6 $\beta_{0,s} + b_{0,r} + (\beta_{F,s} + b_{F,r}) X_F + (\beta_{R,s} + b_{R,r}) X_R$ | $b_{.,r} \sim N(\mu, \sigma)$ | Random / Crossed | Diagonal | 29.1 |
| 7 $\beta_{0,s} + b_{0,r} + (\beta_{F,s} + b_{F,r}) X_F + (\beta_{R,s} + b_{R,r}) X_R$ | $\mathbf{b} \sim MVN(\mu, \Sigma)$ | Random / Crossed | Full | 30.0 |
| 8 $\beta_{0,s} + b_{0,sr} + (\beta_{F,s} + b_{F,sr}) X_F + (\beta_{R,s} + b_{R,sr}) X_R$ | $b_{.,sr} \sim N(\mu, \sigma)$ | Random / Nested | Diagonal | 58.7 |
| 9 $\beta_{0,s} + b_{0,sr} + (\beta_{F,s} + b_{F,sr}) X_F + (\beta_{R,s} + b_{R,sr}) X_R$ | $\mathbf{b} \sim MVN(\mu, \Sigma)$ | Random / Nested | Full | 59.6 |

Parameters were estimated in a Bayesian framework using Hamiltonian-Monte-Carlo (HMC) methods (Neal 2011) and ‘no U-turn sampling’ (NUTS) (Hoffman & Gelman 2014), as implemented in the R package *rstanarm* (Goodrich et al. 2020; Stan Development Team 2020). These sampling techniques are versatile, computationally efficient algorithms for Markov-chain simulation, building on traditional, but less-efficient, approaches such as Gibbs sampling and the Metropolis-Hastings algorithm. For initial model comparison, we ran four chains of 2,000 iterations for each model (due to computational time) and extended this to 12 chains and 4,000 iterations for the selected model. Model selection was performed using approximate leave-one-out cross validation (Vehtari et al. 2017) – this estimates the information-theoretic, relative Kullback-Leibler discrepancy in a Bayesian setting. Model priors were weakly informative, although we found that the time to convergence was reduced by using a slightly narrowed Gaussian prior (*σ* = 0.5) and standardised predictors. We used the R-hat diagnostic (R-hat < 1.01) as well as visual inspection to establish chain convergence (Vehtari et al. 2020). The selected model was validated using posterior predictive checks applied to a summary function based on each of the spatial covariates (Gabry et al. 2019).

## Results

Over a total of 64 survey days, 939 ravens and 1042 roadkill carcasses were observed across all routes and seasons. While roadkill density was generally consistent across survey locations: King (n = 227); Flinders (n = 304); Huon (n = 283); Tasman (n = 228), raven density was much higher on King Island in contrast to the other regions: King (n = 407); Flinders (n = 234); Huon (n = 163); Tasman (n = 135).

We found that models containing random and nested effects (models 8 and 9; Table 1) had the best predictive performance, as estimated by the cross-validation scores (Fig. 3). These two models performed equally well, indicating that the (additional) complexity of including off-diagonal covariance terms (model 9; Table 1) is unjustified; consequently, we infer that the influence of roadkill is independent of the influence of farmland, despite any possible correlation between roadkill locations and habitat type. The simpler models (1-3; Table 1), each of which pool estimates for at least one of the covariates, perform very poorly relative to the best-performing models— where relative performance is calibrated by the estimation uncertainty in the model scores (see error bars in Fig. 3) (Vehtari et al. 2017). ‘Nesting’ of the season-route effects improved the model considerably compared to ‘crossing’ the effects. This suggests that the random effects attributed to each route vary seasonally, above and beyond any direct seasonal effects. The validation plots (Fig.4) suggest that the selected best model (model 8) generates plausible data under posterior model simulation, generating similar trends to the empirical data for the summary function of each spatial covariate.

**Figure 3.**
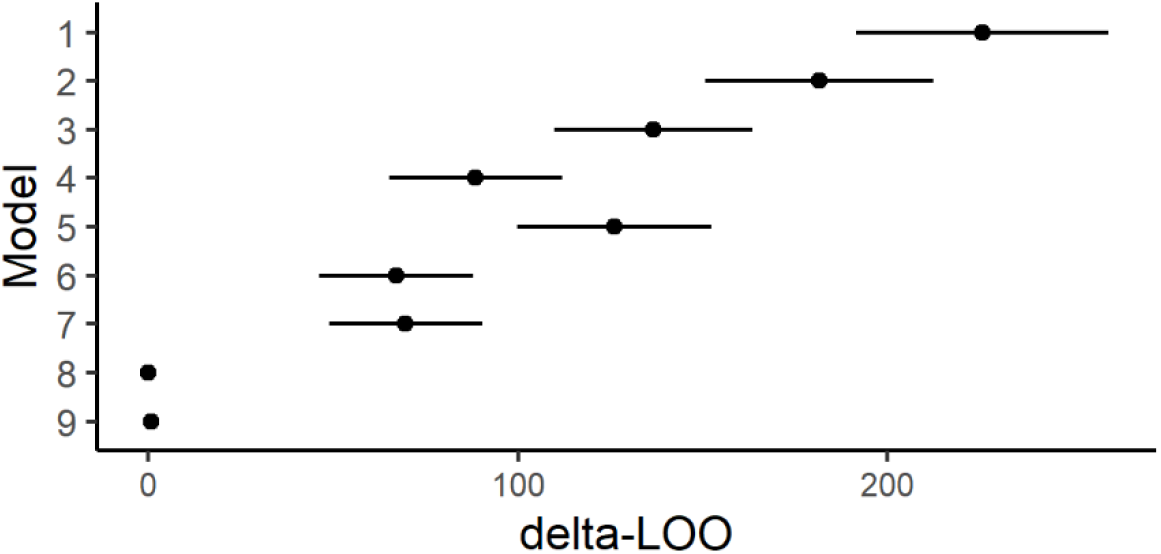
Estimates of model performance using approximate leave-one-out cross validation. The scores ΔLOO_i_= LOO_i_ – LOO_min_ are the differences of the loo estimates of models i = 1,..,9 and that of lowest-scoring model (model 9). The error bars depict one-standard error of the ΔLOO_i_ estimates.

**Figure 4.**
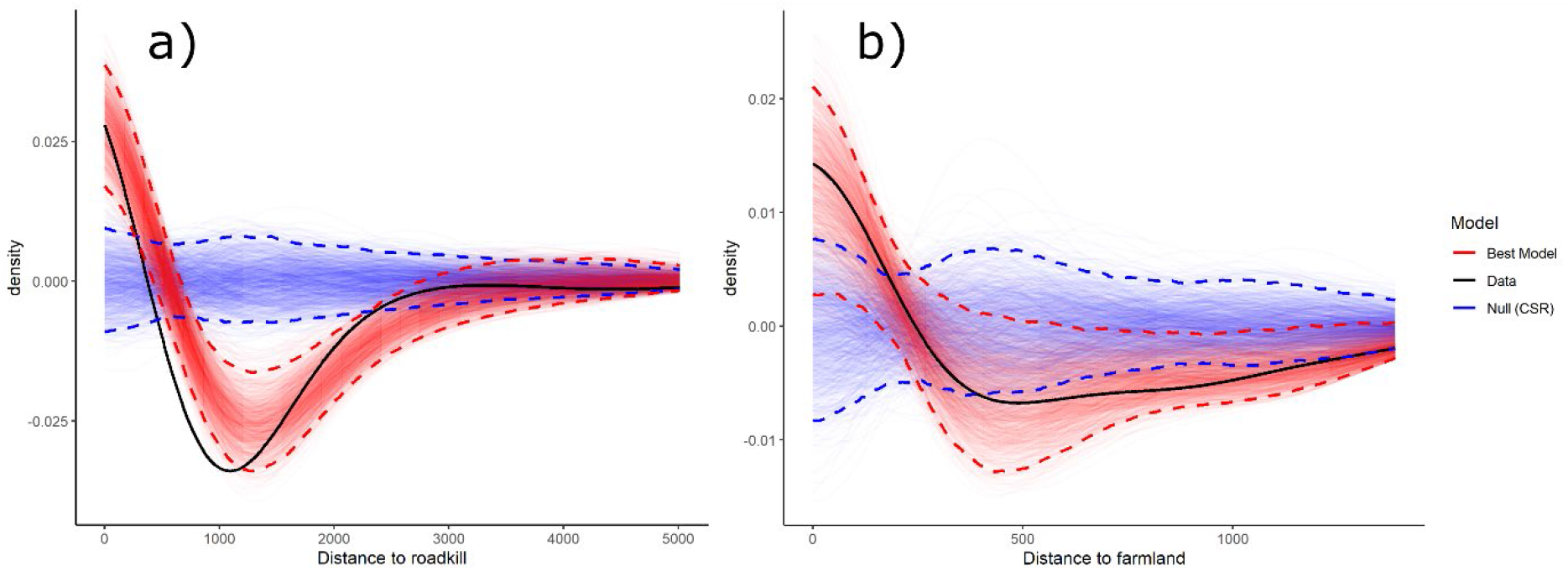
Posterior predictive checks of the selected best model (model 8) for a) distance to roadkill and b) distance to farmland. Generated from 500 posterior draws with density values centred on the mean of the null model.

We found that roadkill was a strong predictor of raven presence along road networks (Fig. 5). On Tasmanian mainland roads, the spatial intensity of ravens was consistently elevated around roadkill across all routes in autumn. The effect of roadkill was amplified on the Bass Strait island routes, where roadside carrion was a predictor of raven presence across the entire year. The effect of distance to farmland on roadside spatial intensity of ravens was weak, but nevertheless consistent across all routes. We observed a weak effect of farmland habitat on raven spatial intensity throughout autumn and, to a lesser extent, winter (Fig. 5). There was no effect of farmland during the summer and spring seasons.

**Figure 5:**
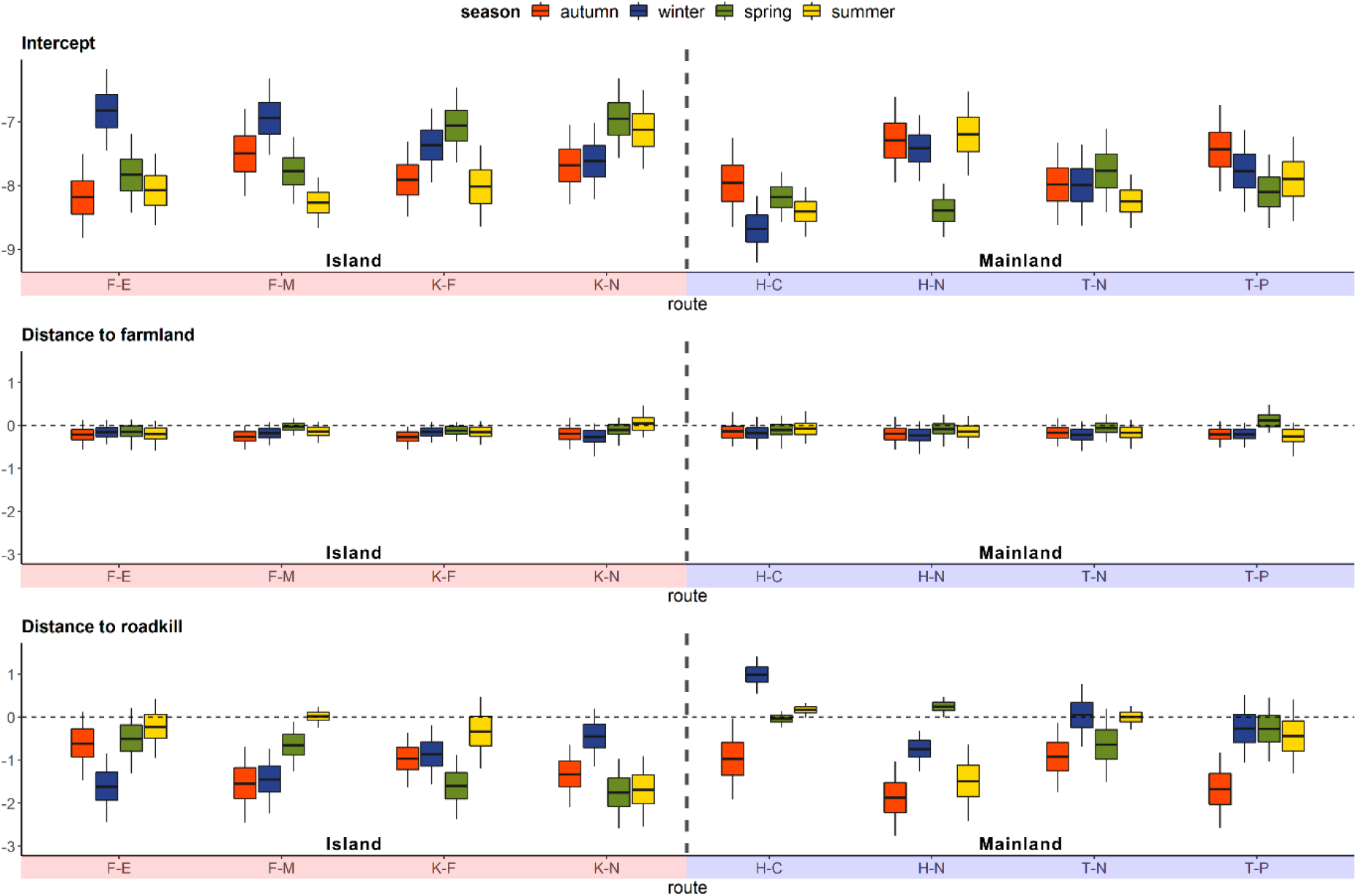
Parameter estimates of raven spatial intensity on the log-linear scale. Distances to both roadkill and farmland have been standardised. The intercept values, which are baseline densities, have been back-transformed to account for the standardisation of the covariates, making the values comparable between routes and seasons. The boxes summarise the marginal posterior distribution for each parameter, displaying the mean and the 50% and 90% credible intervals.

## Discussion

In this study, we used hierarchical Bayesian models to investigate how the spatial pattern of roadside carrion use by a generalist avian scavenger varies across habitat and season, and in response to the (human-mediated) presence or absence of competitor mammalian scavengers. We found that roadkill was a strong predictor of raven presence along roadsides. This roadkill effect was elevated on the Bass Strait islands, where roadside carrion was a predictor of raven presence across the entire year. In contrast, ravens were only highly correlated with roadkill in the autumn months on road networks on the Tasmanian mainland, where they face competition for carcass access from a high diversity of scavenging mammalian predators. We observed a weak effect of farmland habitat on raven presence across all routes during autumn and, to a lesser extent, winter.

The roadkill effect was amplified on the island routes where roadside carrion was a predictor of raven presence across the entire year, except for summer on some routes (Fig. 5). The Bass Strait islands, and King Island in particular, were heavily modified for agriculture following European occupation in the 19^th^ Century (Threatened Species Section 2012). This loss of habitat combined with extensive hunting by Europeans has resulted in the extirpation of several predators on the islands. Both eastern (*Dasyurus viverrinus*) and spotted-tailed quolls (*Dasyurus maculatus*) were hunted to extinction on King and Flinders Islands and a major avian scavenger, the wedge-tailed eagle (*Aquila audax*), is now a rare visitor to King Island (Peacock et al. 2018; Fielding, Buettel & Brook 2020). Remains of the Tasmanian devil (*Sarcophilus harrisii*) have been found in cave deposits on Flinders Island, dated at roughly 4,000 years ago, but it is unclear whether they were present when Europeans arrived, and there is no fossil evidence of the species on King Island (Hope 1973).

The loss of these carnivores from the Bass Strait islands could mean that the remaining scavengers, such as ravens, have less competition for resources and subsequently greater access to carrion. In north-east mainland Tasmania, where the Tasmanian devil has declined due to devil-facial-tumour disease, research has found that forest ravens, feral cats and spotted-tailed quolls all had greater access to carrion, foraging for up to four times as long compared to regions where devils are still abundant (Cunningham et al. 2018). As the spotted-tailed quoll is also absent from the Bass Strait islands due to recent extirpation, forest ravens and feral cats on the islands would have an even greater access to carrion (Peacock et al. 2018). Competition for carrion is known to increase in the warmer months, as carcasses decompose more quickly due to heightened activity by microbes and detritivorous invertebrates (Turner et al. 2017). With the absence of other scavengers on the islands this competition would be reduced, allowing the surviving scavengers to make use of these nutrient-rich resources throughout the year, even when other resources are seasonally available.

Increased scavenging opportunities opened by the loss of apex predators may lead to an increased abundance of the surviving scavengers, affecting ecosystem structure and function (Allen et al. 2014; O’Bryan et al. 2019) In south-eastern Spain, red foxes (*Vulpes vulpes*) were observed to be in higher abundance in areas where the absence of vultures (*Gyps* spp.) increased carrion availability (Morales-Reyes et al. 2017). Cunningham et al. (2018) found that feral cats and forest ravens have increased in abundance where Tasmanian devils have experienced disease-induced declines. On the Bass Strait islands, where the native mammalian apex predators have been extirpated, feral cat and forest raven populations could also be increasing. During the study, we observed a higher number of forest ravens on King Island compared to the other study regions, and anecdotal evidence suggests that forest-raven populations on the island have increased over the last 70 years (Threatened Species Section 2012). The lack of native predators combined with the highly modified landscape of King Island has been suspected as the reasons for increased abundance of forest ravens on the island (Donaghey 2003). Here we have provided evidence for the mechanism underlying this increase—enhanced opportunities for food from roadkill. In Tasmania, past research found that forest raven abundance had no observable impact on the abundance of other bird species (Fielding, Buettel, Nguyen, et al. 2020). However, these impacts may be present on the Bass Strait Islands where ravens are in higher abundance, potentially leading to heightened depredation on the nestlings and eggs of other birds, impacting the survival of other species (Read & Wilson 2004; Madden et al. 2015; Webb et al. 2016). Conversely, ravens on the islands may be preying on nestlings and small birds less due to their uninterrupted access to roadkill across the entire year – this interaction requires further investigation.

The effect of roadkill on raven presence was only consistent across all routes, both island and mainland, during the autumn surveys (Fig. 5). During the autumn months, there are fewer alternative food sources, such as lizards, insects, or fruit, for facultative scavengers like the forest raven (Rowley & Vestjens 1973). Additionally, as autumn is outside of the breeding season, there are fewer juvenile animals and therefore less opportunities for easy prey (Higgins et al. 2006).

Therefore, scavengers may be more dependent on carrion, such as roadkill, for food during the autumn months (Hobday & Minstrell 2008; Fielding, Buettel, Nguyen, et al. 2020).

Interestingly, there was a weak effect of farmland on the presence of ravens during autumn across all routes (Fig. 5). During autumn, farmers will sow seeds for spring growth, including beef farmers that will sow oats and rye for winter stock food. As ravens are omnivorous, they may be utilising the soil disturbance during sowing to access grubs and worms (Rowley & Vestjens 1973; Higgins et al. 2006). Some farmers will also lamb or calf during the autumn instead of spring. Local farmers on Flinders Island report that ravens will follow lamb breeding schedules across the island, moving from paddock to paddock as stock start to give birth. The design of the study meant that we could not detect this paddock migration, but it warrants further investigation. Additionally, future research could investigate and contrast how ravens use carcasses in paddocks, such as culled macropods, compared to roadside carrion.

Model validation of the effect of distance to roadkill on raven density showed that the (posterior) predictions of the selected best model followed the general shape of the observed data, but slightly overestimated the range of the effect (Fig. 4a). This result suggests that there may be additional predictors influencing the presence of forest ravens along roadsides that were not measured in this study and should be investigated further. While point-pattern methods have been used for multi-dimensional data (e.g., two-dimensional vegetation plots), they have been underutilised for one-dimensional analysis (Baddeley et al. 2020). Our Bayesian approach to parameter estimation of Poisson point-process models facilitates the inclusion of hierarchical structures and subsequent model comparison. Alternative approaches based on maximum-likelihood estimation should also be possible, although algorithm convergence and model comparison can be problematic when hierarchical effects are nested or crossed and candidate models have differing effect structures (Bolker et al. 2009). The approach we have developed herein to analyse the roadkill problem has broad applicability within ecological research and could be used with other one-dimensional datasets, such as survey transects or animal tracks.

Our study provides evidence that the loss of native predators is beneficial for generalist scavengers, enabling year-round use of roadside carrion. This may trigger trophic shifts, leading to increased populations of certain species and resulting in structural changes and potentially unstable imbalances within the ecosystem (Allen et al. 2015; Buechley & Şekercioğlu 2016). However, these population shifts, and the associated consequences, require further investigation. Regardless, the results of this study highlight the importance of conservation intervention in these circumstances, such as the targeted management of roadside carrion (e.g., removal of carcasses or vehicle speed reduction) and/or the reintroduction of the recently extirpated native scavengers (Fielding, Buettel & Brook 2020). As we are faced with the escalating threat of further extinctions across the world, our study highlights the significance of conserving scavenger diversity within an ecosystem.

## Acknowledgements

The authors would like to acknowledge the Palawa peoples of lutruwita, the traditional custodians of the lands on which this work was completed. We gratefully acknowledge those who assisted with data collection – Hanh Nguyen, Alexander Gordon, Lucas Sayer, Patrick Gee, Isaac Gee, Hans Ammitzboll, Glenn Goessing, Victoria Paine, Margaret Batey & Lizzie Cambra.

## Authors’ contributions

M.W.F., B.W.B. and J.C.B. designed the study, developed the field methodology and collected the data; L.Y. developed the analysis; L.Y. and M.W.F. analysed the data and wrote the manuscript. All authors contributed significantly to the drafts and gave final approval for publication. The authors claim no conflicts of interest.

## Data Availability Statement

Data available from the figshare repository https://doi.org/10.6084/m9.figshare.14036807 (Fielding et al. 2021). Code available from the Zenodo Repository: http://doi.org/10.5281/zenodo.4549778 (Yates et al. 2021)

